# Effects of different tradititonal and commercial aquafeed on proximate composition and growth performance of Grass carp (*Ctenopharyngodon idella*) reared in the semi-intensive composite culture system

**DOI:** 10.1101/2023.02.20.529235

**Authors:** Talha Zulfiqar, Muhammad Sajjad Sarwar, Habib Ul Hassan, Abdul Shakoor Chaudhry, Muhammad Hafeez-ur-Rehman

**Affiliations:** Department of Zoology, Faculty of Life Sciences, University of Okara, Okara, 56300, Pakistan; Department of Zoology, University of Karachi,75270 Karachi, Pakistan; Agriculture Building Newcastle University Newcastle upon Tyne UK NE1 7RU; Department of Fisheries and Aquaculture, University of Veterinary and Animal Sciences, Lahore, Pakistan

**Author notes:** Kind regards, Habib- Ul- Hassan, Department of Zoology (MRCC), University of Karachi, Karachi-75270, Pakistan.

**Keywords:** Commercial aqufeed, Traditional feed, Growth, Body compostion, Grass carp

## Abstract

The objective of the research was to find the effects of numerous traditional and commercial aqua feed on proximate composition and growth performance of Grass carp (*Ctenopharyngodon idella*) reared in the semi intensive composite culture system. The aqua feeds of various companies (AMG, Supreme, Aqua, Star Floating, Hi-Pro and Punjab feed) used as commercial feed. Farm made feeds were Maize gluten and Rice polish. For confidentiality, these feeds were randomly assign code T1, T2, T3, T4, T5, T6, T7 and T8 which were only known to investigating staffs. There were two replicates for each treatment. In this research, significantly maximum growth was recorded in T3 as compared to other treatments. Lesser weight gain was observed in the T1 (270.30±60.5).The maximum body length (19.25±2.19) was found in T3. Similarly, the minimum body length (5.97±2.94) was seen in T2. FCR ratio (2.36±0.01) was recorded in T3. Simultaneously, FCR (1.86±0.002) was also recorded in T4 that is the perfect ratio for fish farmers. Higher SGR was found in T3 (1.62±0.05). Overall, T4 showed lesser SGR (1.05±0.001). T4 showed the higher Crude protein (28.66±0.24%). T3 showed more fat content level (5.46±0.33%) in the body. These outcomes also supported that enrichment in the dietary amount of protein and lipid level improve lipids content and crude protein in fish body weight. Thus, based on growth performance, survival and proximate composition, it is concluded that T3 & T4 may be recommended for commercial culture of *Ctenopharyngodon idella*.

## 1. Introduction

Aquaculture is the developing as a sustainable source of foodstuff for individuals and also imparts its role in the well-being of human beings[1, 2]. Fish is a richsource of high quality protein, numerous fatty acids as well as micronutrients [3–5]. Fish contribute more to provide food and nutrition value than other basic food contents like vegetables [6]. World population is increasing rapidly and demand of fish and fish products are also increasing. So, maintaining per capita fish consumption can be done by increasing aquaculture production. Grass carp (*Ctenopharyngodon idella*) is herbivorous exotic fish introduced for its food on macrophages and adult grasses. Grass carp eat about 24% of plant and 76% animal food [7]. Grass carp depends upon aquatic plants and weeds for its growth [7, 8]. It feeds on aquatic vegetation and has pharyngeal teeth to control aquatic grasses [9]. Feed consumption level can be increased in aquaculture system by giving fish a good healthy commercial feed that increase its growth[10]. Semi intensive system in ponds by fertilizers and commercial feeding is means of production low cost fish in emerging countries [11–15]. For better growth of aquaculture it is necessary to sustain the means of feed and fertilizers [16]. Dietary protein is essential for suitable working, growth and make over of different body protein [17, 18]. However, fats and carbohydrates are also necessary for fish energy utilization and these also help to improve consumption of dietary protein [19, 20]. Culture of fast growing major carps such as *Ct. idella* helps to eliminate the problems of protein deficiency [21, 22]. The goal of semi intensive fish culturing is to produce huge amount of fish in a limited time period [23, 24]. Utilization of commercial feed is very important for the development of semi intensive composite culture of Grass carp with other major carps[25, 26]. Fish feed components should provide balanced minerals, amino acids and energy[10].

## 2. Materials and methods

### 2.1 Study area

The study was carried out for proximate analysis of commercial aquafeed and fish feed ingredients (Corn Gluten, Rice Polish) purchased by visiting various commercial feed mills and numerous fish ponds in different regions of the Punjab, Pakistan by using the facilities of research laboratories at Department of Fisheries and Aquaculture, University of Veterinary and Animal Sciences (UVAS), Ravi Campus, Pattoki, Punjab, Pakistan.

### 2.2 Experimental design

Two types of farm made and Six types of commercial aquafeed containing up to 20 to 30% CP provided by Pvt. Ltd. given in treated ponds at 2% feed of fish wet body weight two times per day for six months. The aquafeed of various companies (Supreme Feed, AMG Feed, Aqua feed, Star Floating feed, Hi-Pro feed and Punjab feed) were used as commercial feed. Sixteen ponds were treated with this feed. Farm made aqua feed were Corn gluten and Rice polish. Treatement design for confidentialy these feed were randomly give code T1, T2, T3, T4, T5, T6, T7 and T8 which were only known to investigating staffs. There were two replicates for each treatment. The ponds were filled up by water up to 5 feet. Inorganic and organic manures were used to increase the fertility of ponds.

### 2.3 Growth performance and analysis

Total body length and wet body weight of experimental fish were measured at the time of initial stocking. While during research, fish were sampled on fortnightly basis to evaluate the effect of the feed on body length and total body weight. Net weight gain, feed conversion ratio (FCR) and Specific growth rate (SGR were calculated according to give formula.

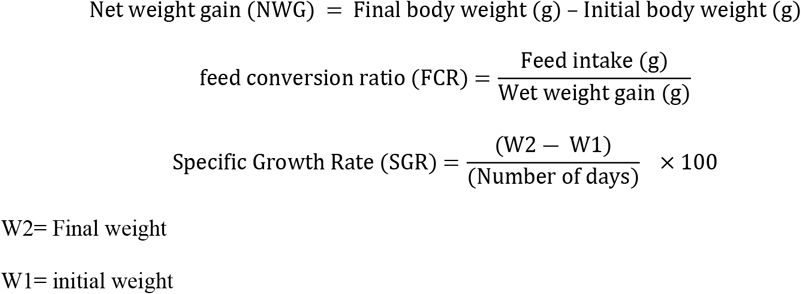

### 2.4 Proximate anaylsis

#### Moisture contents

(4-5g) sample taken enclosed with smooth aluminium and dehydrated at 100°C to constant weight in furnace fixed with measured aeration. The weight reduced was note down as moisture.

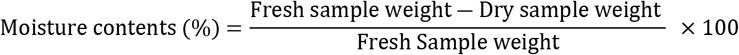

#### Crude protein

1g Sample was mixed with 0.7g mercuric oxide as a catalyst,10g potassium sulphate then processed in a long Kjeldhal flasks for exactly 2 hours (one hour after contents are clear) with concentrated 20ml sulphuric acid at an inclined angle after the addition of 25ml sulphide solution,80ml of 40% NaOH,90ml distilled water and were placed in the apparatus two layers were formed while oriented the flask then attached to condenser unit, collected ammonia in 50ml of solution of boric acid by using few drop of methyl red as an indicator and collected ammonia for almost 2 minutes later the colour changed from pink to golden yellow. Then solution of ammonia in boric acid was titrated with 0.1N HCl. The volume of HCl used was recorded and nitrogen (%) calculated as below:

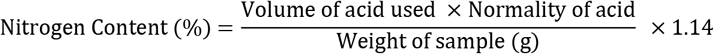

Content of crude protein was calculated by:

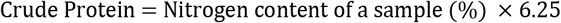

#### Crude fat

2g samples of dry feed were shifted to Soxhlet thimble then the thimble remained retained into a Soxhlet apparatus. Put a dehydrated, tared solvent flask in place below, after adding 25ml ether solvent attached to condenser. Maintain warming ratio to provide 2-3 drops/second condensation rate and extracted for 6 hours. Then removed thimble and regained ether by the apparatus. The elimination of ether was done on a hot water bath and the flask was dehydrated at 105°C for 30 minutes and then cooled in a desiccator and weighed.

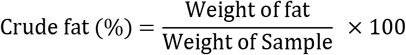

#### Crude fibre

2g feed samples of fat free weighed into a 600ml beaker after the adding of 200ml hot sulphuric acid. Under condenser placed the beaker and for 30 minutes carried to boiling, maintain volume using distilled water and to rinse down particles sticking to the edges. By using Whatman paper No. 541 filtered in a Buchner funnel by suction and washed well with hot water. Filtrate was moved back to the beaker after the addition of 20ml warm sodium hydroxide solution. Placed in condenser then for 1 minute was carried to boiling. Approximately 30 minutes afterward boiling, filtered by a porous crucible and was rinsed through hot water, 1% HCl and once more by hot water. Rinsed two times by alcohol, at 100°C dehydrated overnight ash for 3 hours at 500°C, cooled then weighed.

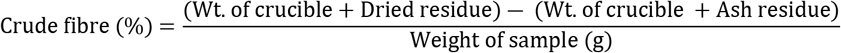

#### Ash content

2g feed samples was dehydrated tarred porcelain dish then placed in a furnace at 600 °C for 6 hours. Kept in desiccator for cooling and then weighed.

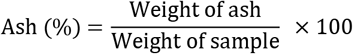

#### Statistical analysis

The impact of commercial aqua feed and farm made feed on proximate composition and growth rate were evaluated by one-way ANOVA techniques. The General Linear Model process applied through SAS software (version 9.1). Duncan’s Multiple Range Test (DMRT) was used to find out the Means of Significant of treatments.

## 3. RESULTS

The average initial and final body weight (g) as well as net gain in body weight (g) of *Ctenopharyngodon idella* are mentioned in table 1 that were fed with different feeds (commercial and traditional). The treatment 1 and 2 were served with traditional aqua feeds and treatment 3,4,5,6,7,and 8 were treated with commercial feed. The average initial body weight of *Ctenopharyngodon idella* of treatment 1 and 2 were 266.00±46.18 g and 268.00±46.09 g, respectively. Similarly, the average final body weight of treatment 1 and treatment 2 were 631.95±153.08 g and 537.30±72.60 g, respectively. The net weight gain in treatment 1 was 365.95 g and net weight gain of treatment 2 was 269.30 g.

**Table 1.** Proximate analysis of farm-made and commercial aquafeeds.

| **Treatments** | **T1** | **T2** | **T3** | **T4** | **T5** | **T6** | **T7** | **T8** |
| --- | --- | --- | --- | --- | --- | --- | --- | --- |
| **Crude Protein (%)** | 14.64 ± 0.09 | 25.66 ± 0.08 | 22.20±1.13 | 24.23±1.47 | 22.40 ±3.08 | 25.20 ±1.67 | 24.328±2.68 | 22.54±1.8 |
| **Crude Fats (%)** | 3.08 ± 0.09 | 0.47 ± 0.04 | 4.44±2.69 | 4.44 ±0.17 | 6.63 ±2.15 | 3.12 ±1.2 | 13.449±0.96 | 14.47±2.02 |
| **Ash (%)** | 6.24 ± 0.07 | 9.77 ± 0.02 | 23.6±0.50 | 25.7±1.46 | 27.8 ±1.9 | 12.51±3.76 | 11.464±2.38 | 13.35±1.38 |
| **Crude Fiber(%)** | 7.21 ±1.23 | 9.10 ±1.01 | 8.17 ±1.37 | 7.38 ±0.61 | 6.97±1.31 | 9.45±1.18 | 9.97±0.243 | 8.78±1.05 |
| **Dry matter (%)** | 95.1± 0.23 | 90.3± 0.21 | 93.1± 0.04 | 94.8 ± 0.06 | 90.2± 0.13 | 94.4± 0.09 | 90.5± 0.08 | 93.4± 0.14 |
| **Moisture (%)** | 4.86 ± 0.07 | 12.21 ± 0.05 | 6.9±3.05 | 5.2±0.34 | 8.3±1.24 | 7.8 ±5.57 | 12.622±1.35 | 14.34±0.66 |

The average initial body weight of treatment 3 was 272.50±50.74 g and the average initial body weight of treatment 4 was 279.00±48.67 g. Similarly, average final body weight in treatment 3 and treatment 4 were 766.40±108.42 g and 646.15±118.54 g, respectively. The net body weight gain of treatment 3 and treatment 4 were 493.90 g and 367.15 g, respectively.

The average initial body weight of treatment 5 was 339.00±74.38 g and the average initial body weight of treatment 6 was 336.00±76.98 g. Similarly, the average final body weight of treatment 5 and treatment 6 were 870.90±204.05 g and 721.20±222.63 g, respectively. The net body weight gain of treatment 5 and treatment 6 were 531.90 g and 385.2 g, respectively.

The average initial body weight of treatment 7 and treatment 8 were 186.00±25.53 g and 184.50±23.67 g, respectively. Similarly, the average final body weight of treatment 7 and treatment 8 were 698.65±127.77 g and 559.60±67.73 g, respectively. Net weight gain of treatment 7 was 512.65 g and the net gain weight of treatment 8 is 375.10 g.

**Table 2.** The average initial body weight (g) and length (cm), average final body weight and length and gain in body weight and length, SGR and FCR of Grass carp (*Ctenopharyngodon idella*) fed with different feeds (farm made feed and commercial aqua feed).

| Parameters | T1 | T2 | T3 | T4 | T5 | T6 | T7 | T8 |
| --- | --- | --- | --- | --- | --- | --- | --- | --- |
| **Initial weight(g)** | 266.00±46.18c | 268.00±46.09c | 272.50±50.74c | 279.00±48.67c | 339.00±74.38c | 336.00±76.98c | 186.00±25.53c | 184.50±23.67c |
| **Final weight(g)** | 631.95±153.08a | 537.30±72.60b | 766.40±108.42a | 646.15±118.54b | 870.90±204.05a | 721.20±222.63b | 698.65±127.77a | 559.60±67.73b |
| **Net weight gain(g)** | 365.95 | 269.30 | 493.90 | 367.15 | 531.90 | 385.2 | 512.65 | 375.10 |
| **Initial length(cm)** | 26.00±2.23c | 25.00±2.46c | 28.50±3.48c | 29.50±3.17c | 27.00±3.44b | 26.50±3.01b | 18.50±2.21c | 18.50±2.21c |
| **Final length(cm)** | 35.85±2.36a | 33.35±1.63b | 41.00±3.24a | 35.47±2.70b | 38.20±1.85a | 36.92±1.83a | 37.75±2.17a | 33.50±1.79b |
| **Net length gain(g)** | 9.85 | 8.35 | 12.5 | 5.97 | 11.2 | 10.42 | 19.25 | 15 |
| **FCR** | 2.69a | 2.82a | 2.64a | 3.02a | 1.86a | 3.39a | 2.36a | 2.53a |
| **SGR** | 1.14a | 1.23a | 1.26a | 1.21a | 1.05a | 1.39a | 1.62a | 1.36a |

Means with similar letters are not significantly different.

**Table 3.** The average values of proximate analysis of Grass carp (*Ctenopharyngodon idella*) that were fed by farm made feed & commercial aqua feed.

| Parameters | T1 | T2 | T3 | T4 | T5 | T6 | T7 | T8 |
| --- | --- | --- | --- | --- | --- | --- | --- | --- |
| **Crude protein (%)** | 28.66a | 21.97a | 24.45c | 17.62b | 25.80a | 22.40b | 24.65a | 19.01a |
| **Crude fat (%)** | 3.99a | 2.36b | 2.40a | 1.30c | 5.46b | 3.64a | 2.13b | 1.80a |
| **Dry matter (%)** | 22.90a | 21.67a | 19.45a | 21.98a | 22.67a | 21.45a | 22.81a | 22.64a |
| **Ash (%)** | 2.85a | 2.65a | 1.98b | 1.67c | 2.34a | 2.07c | 2.16a | 2.09b |

Means with same letters are not significantly different.

**Table 4.** Average values of physic chemical parameters of pond water under the influence of different traditional and commercial aqua feed.

| Parameters | T1 | T2 | T3 | T4 | T5 | T6 | T7 | T8 |
| --- | --- | --- | --- | --- | --- | --- | --- | --- |
| **Temperature** | 25.8±0.35<sup>a</sup> | 26.1±0.25<sup>a</sup> | 26.7±0.34<sup>a</sup> | 25.3±0.14<sup>a</sup> | 24.7±0.61<sup>a</sup> | 27.2±0.15<sup>a</sup> | 25.1±0.14<sup>a</sup> | 26.3±0.32<sup>a</sup> |
| **pH** | 7.92±0.34<sup>a</sup> | 8.19±0.11<sup>a</sup> | 7.89±0.68<sup>a</sup> | 8.05±0.27<sup>a</sup> | 8.02±0.31<sup>a</sup> | 7.90±0.14<sup>a</sup> | 7.79±0.61<sup>a</sup> | 8.12±0.35<sup>a</sup> |
| **DO** | 6.98±0.58<sup>a</sup> | 7.15±0.22<sup>a</sup> | 7.67±0.45<sup>a</sup> | 7.49±0.78<sup>a</sup> | 7.55±0.32<sup>a</sup> | 6.35±0.18<sup>a</sup> | 7.54±0.43<sup>a</sup> | 6.76±0.27<sup>a</sup> |
| **TDS (ppm)** | 680±0.76<sup>a</sup> | 775±0.49<sup>a</sup> | 734±0.24<sup>a</sup> | 699±0.33<sup>a</sup> | 795±0.65<sup>a</sup> | 780±0.51<sup>a</sup> | 699±0.42<sup>a</sup> | 791±0.22<sup>a</sup> |

Means with same letters are not significantly different.

**Figure 1.**
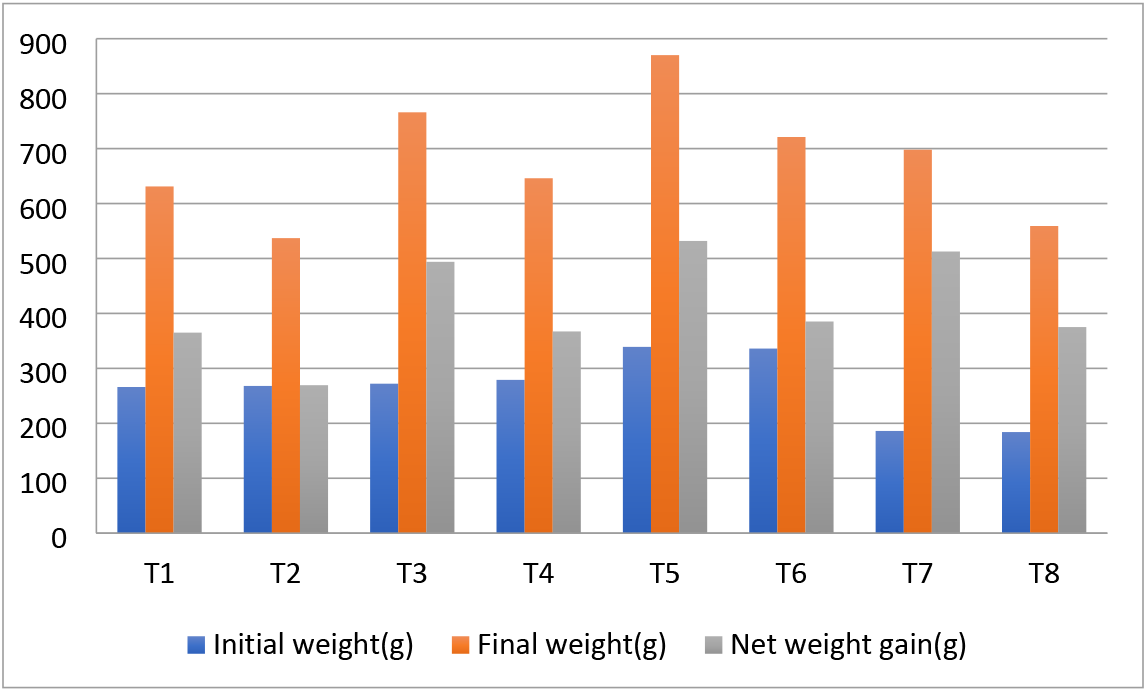
The average initial body weight (g), average final body weight and net gain in body weight of Grass carp (*Ctenopharyngodon idella*) fed with different feeds.

**Figure 2.**
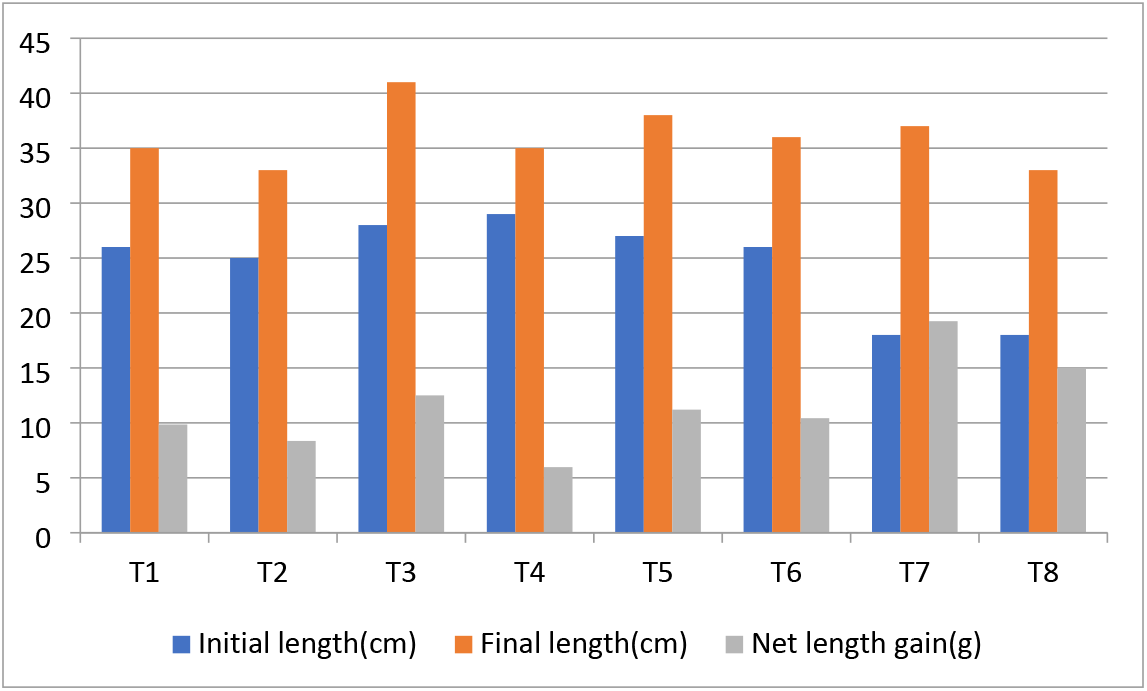
The average initial body length (cm), average final body length and net gain in body length of Grass carp (*Ctenopharyngodon idella*) fed with different feeds.

**Figure 3.**
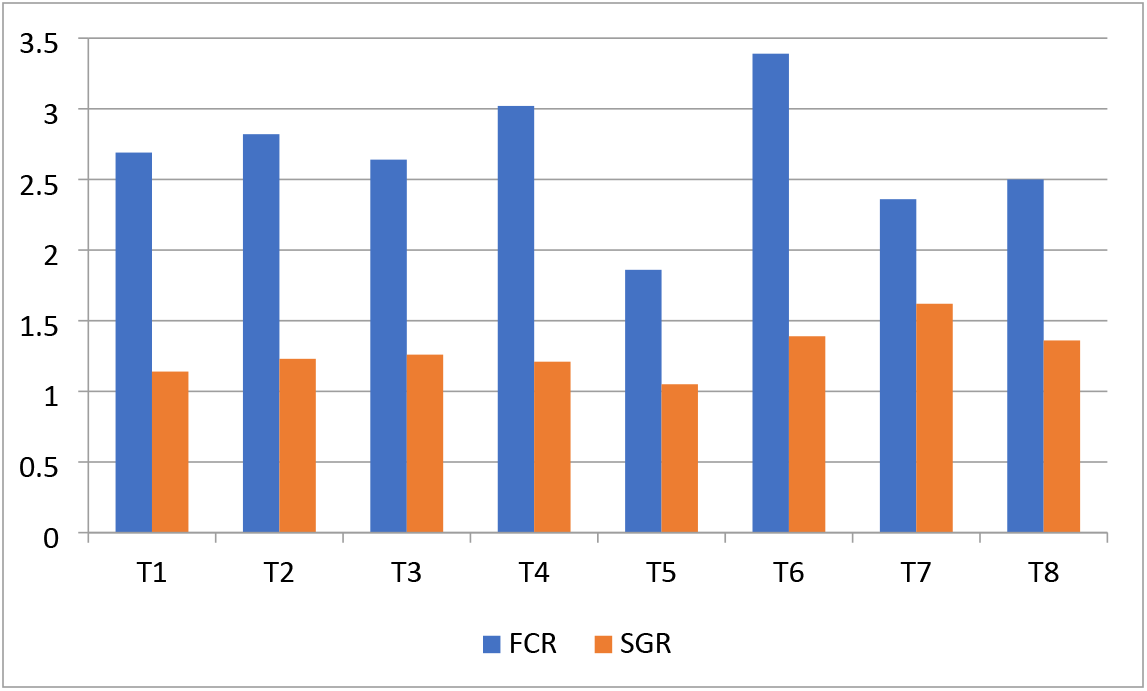
The feed conversion ratio (FCR) Specific growth rate (SGR) of Grass carp (*Ctenopharyngodon idella*) fed with different feeds.

**Figure 4.**
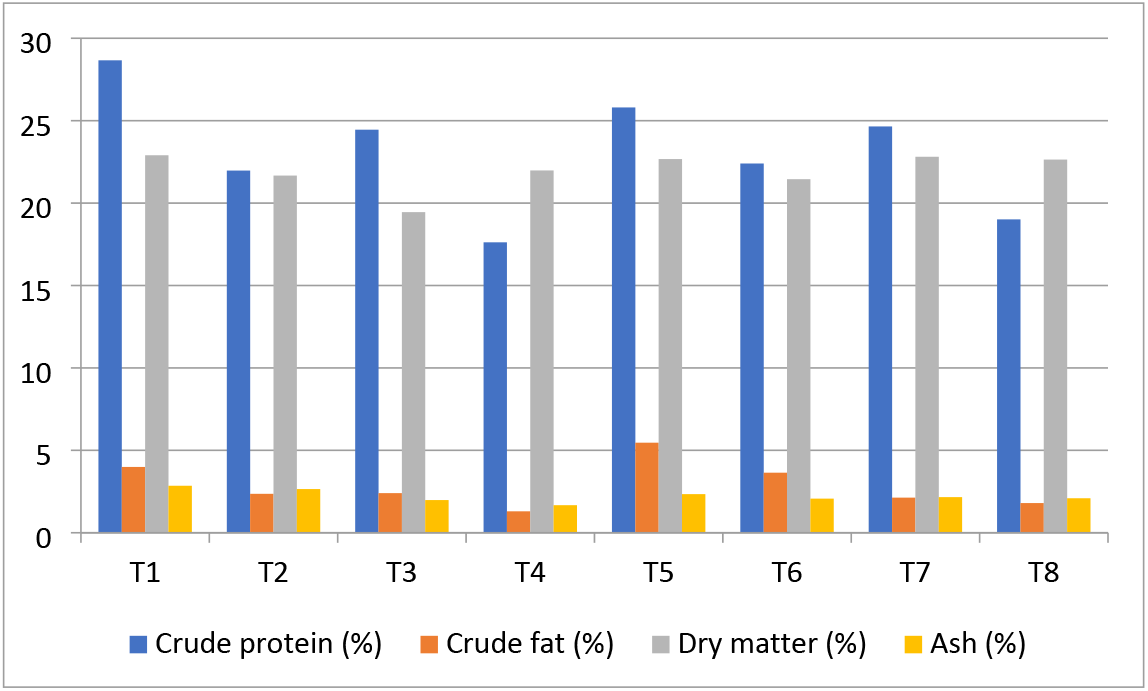
The average values of proximate analysis of Grass carp (*Ctenopharyngodon idella*) that were fed by farm made feed & commercial aqua feed.

The average initial and final body length (g) as well as net gain in body length (cm) of *Ctenopharyngodon idella* are mentioned in table 2 that are fed with different feed (commercial and traditional). The average initial body length of treatment 1 and treatment 2 were 26.00±2.23 cm and 25.00±2.46 cm, respectively. Similarly, average final body length of treatment 1 and 2 remained 35.85±2.36 cm and 33.35±1.63 cm, respectively. The net length gain of treatment 1 was 9.85 cm and the net weight gain of treatment 2 was 8.35 cm. The average initial body length of treatment 3 and 4 remained 28.50±3.48 cm and 29.50±3.17 cm, respectively. Similarly, average final body length of treatment 3 and 4 were 41.00±3.24 cm and 35.47±2.70 cm, respectively. The net length gain of treatment 3 was 12.5 cm and net weight gain of treatment 4 was 5.97 cm. The average initial body length of treatment 5 was 27.00±3.44 cm and the average initial body length of treatment 6 was 26.50±3.01 cm. Similarly, the average final body length of treatment 5 was 38.20±1.85 cm and the average final body length of treatment 6 was 36.92±1.83 cm. The net length gain of treatment 4=5 and 6 was 11.2 cm and 10.42 cm, respectively. The average initial body length of treatment 7 and 8 was 18.50±2.21 cm and 18.50±2.21 cm, respectively. Similarly, the average final body length of treatment 7 and treatment 8 was 37.75±2.17 cm and 33.50±1.79 cm, respectively. The net length gain of treatment 7 was 19.25 cm and the net weight gain of treatment 8 was 15 cm. The average value of feed conversion ratio (FCR) of treatment 1 and 2 were 2.69 and 2.82, respectively. The average specific growth rate (SGR) of treatment 1 was 1.14 and the specific growth rate (SGR) of treatment 2 was 1.23. The average feed conversion ratio (FCR) of treatment 3 was 2.64. Similarly, the average feed conversion ratio (FCR) of treatment 4 was 3.02. The average value specific growth rate (SGR) of treatment 3 was 1.26 and the value specific growth rate (SGR) of intreatment 4 was 1.21.

The average feed conversion ratio (FCR) of treatment 5 and 6 were 1.86 and 3.39, respectively. The average specific growth rate (SGR) of treatment 5 was 1.05 and specific growth rate (SGR) of treatment 6 was 1.39. The average feed conversion ratio (FCR) of treatment 7 was 2.36. Similarly, the average feed conversion ratio (FCR) of treatment 8 was 2.53. The average specific growth rate (SGR) of treatment 8 was 1.62 and specific growth rate (SGR) of treatment 8 was 1.36.

The average percentage of crude protein (CP) level of treatment 1 was recorded 28.66 % and the average percentage level of crude protein (CP) of treatment 2 was recorded 21.97 %. Similarly, the average value of crude protein (CP) level of treatment 3 and 4 were calculated 24.45% and 17.62%, respectively. The average value crude protein (CP) level of treatment 5 was recorded 25.80% and the average value crude protein (CP) level of treatment 6 was calculated 22.40%. Similarly, the average value crude protein (CP) level of treatment 7 and 8 were recorded 24.65 % and 19.01 %, respectively.

The average percentage of crude fat (CF) level of treatment 1 was recorded 3.99 % and the average percentage of crude fat (CF) level of treatment 2 was recorded 2.36 %. Similarly, the average percentage level of crude fat (CF) of treatment 3 and treatment 4 was calculated 2.40a % and 1.30 %, respectively. The average value crude fat (CF) level of treatment 5 was recorded 5.46 % and the average value crude fat (CF) level of treatment 6 was calculated 3.64 %. Similarly, the average percentage of crude fat (CF) level of treatment 7 and were recorded 2.13 % and 1.80 %, respectively.

The average percentage of dry matter level of treatment 1 was recorded 22.90 % and the average percentage of dry matter level of treatment 2 was recorded 21.67 %. Similarly, the average percentage of dry matter level of treatment 3 and 4 were calculated 19.45 % and 21.98 %, respectively. The average percentage of dry matter level of treatment 5 was recorded 22.67 % and the average percentage level of dry matter of treatment 6 was calculated 21.45 %. Similarly, the average percentage of dry matter level of treatment 7 and 8 were recorded 22.81 % and 22.64 %, respectively. The average percentage of ash level of treatment 1 was recorded 2.85 % and the average percentage of ash level of treatment 2 was recorded 2.65 %. Similarly, the average percentage of ash level of treatment 3 and 4 were calculated 1.98 % and 1.67 %, respectively. The average percentage of dry ash level of treatment 5 was recorded 1.67 % and the average percentage level of ash in treatment 6 was calculated 2.07 %. Similarly, the average percentage of ash level of treatment 7 and treatment 8 was recorded 2.16 % and 2.09 %, respectivel.

## 4. Discussion

Aim of aquaculture is to provide maximum growth and better yield with optimum qualitative characteristic of meat [27]. Fish growth is recognized as increased in total net weight and length of fish by using feeding regime [28]. In this research, significantly maximum growth was found in T3 as compare to other treatments. In T3 the weight gain was lower (512.65±78.61) as compared to T4. Overall, In the T1 lesser net weight gain wer recorded (270.30±60.5). There were significant variation in body length fed with commercial aquafeed and farm made feeds. The significantly maximum average body length 19.25±2.19 was found in treatment 3 fed with commercial aquafeed containing crude protein (CP) level of 26.705±1.13. Similarly, the minimum average body length 5.97±2.94 was seen in T2 farm made feeds with (CP) level of 25.66 ± 0.08. FCR ratio (2.36±0.01) was also recorded in T3. These outcomes were compared by the findings *[29, 30]*. Simultaneously, Best ratio of FCR was recorded in T4 (1.86±0.002). Significantly higher SGR was recorded in T3 (1.62±0.05). As compared to T1 and T7 higher SGR was recoreded in T8 (1.39±0.07) and SGR (1.36±0.039) of T7 higher than T1. Overall, lesser SGR (1.05±0.001) was recorded in T4. These results match with the findings [30]. In our results, T4 treatment groups showed the higher Crude protein (28.66±0.24%) as compared to other treatments by comparing with [27]. These results also evidenced that dietry protein level in the fish body can be enahnced by increasing protein level of aquafeed [17, 31, 32]. Aquafeed with highest lipid content showed the most elevated crude lipids in the fish body [33]. Higher fat content(5.46±0.33%) was recorded in T3 as compare to other treatments.

Physico-chemical parameters like Temperature, pH, alkalinity, phosphate, Nitrate, Dissolved oxygen (DO) and Hardness of water in ponds for different feed treatments did not show any significant variations [27, 30]. The recorded values were within the optimum range.

## Conclusion

During the present study it was found that the average increase in body weight of treatments fed with commercial feeds was greater than treatments fed with traditional feeds. Similarly, the average increase in body length and FCR and SGR of treatments fed with traditional feeds was not recorded as good as of treatments fed with commercial feeds. It was also calculated that the physico chemical parameters of treatment fed with commercial feeds was recorded in good range as compared to treatment fed with traditional feeds. The results of proximate analysis of treatments that fed with commercial feeds were also good as compared to treatments that fed with traditional feeds.

